# Utilization of tissue ploidy level variation in *de novo* transcriptome assembly of *Pinus sylvestris*

**DOI:** 10.1101/495689

**Authors:** Dario I. Ojeda, Tiina Mattila, Tom Ruttink, Sonja Kujala, Katri Kärkkäinen, Jukka-Pekka Verta, Tanja Pyhäjärvi

## Abstract

Compared to angiosperms, gymnosperms lag behind in the availability of assembled and annotated genomes. Most genomic analyses in gymnosperms, especially conifer tree species, rely on the use of *de novo* assembled transcriptomes. However, the level of allelic redundancy and transcript fragmentation in these assembled transcriptomes, and their effect on downstream applications have not been fully investigated. Here, we assessed three assembly strategies, including the utility of haploid (megagametophyte) tissue during *de novo* assembly as single-allele guides, for six individuals and five different tissues in *Pinus sylvestris*. We then contrasted haploid and diploid tissue genotype calls obtained from the assembled transcriptomes to evaluate the extent of paralog mapping. The use of the haploid tissue during assembly increased its completeness without reducing the number of assembled transcripts. Our results suggest that current strategies that rely on available genomic resources as guidance to minimize allelic redundancy are less effective than the application of strategies that cluster redundant assembled transcripts. The strategy yielding the lowest levels of allelic redundancy among the assembled transcriptomes assessed here was the generation of SuperTranscripts with Lace followed by CD-HIT clustering. However, we still observed some levels of heterozygosity (multiple gene fragments per transcript reflecting allelic redundancy) in this assembled transcriptome on the haploid tissue, indicating that further filtering is required before using these assemblies for downstream applications. We discuss the influence of allelic redundancy when these reference transcriptomes are used to select regions for probe design of exome capture baits and for estimation of population genetic diversity.

## 1. INTRODUCTION

Coniferous trees are a dominant member of boreal forests worldwide. From an ecological, genetic, and evolutionary point of view, they represent an interesting group for comparative analyses to other seed plant groups. Due to their phylogenetic position (Z. Li et al., 2015), variation in genome size, highly repetitive structure and organization rich in pseudogenes (Wan et al., 2018), they can serve as a useful contrasting group for comparative genomic analyses with angiosperms. However, the limited availability of genomic resources, particularly whole genome reference sequences (Birol et al., 2013; Neale et al., 2014, 2017; Nystedt et al., 2013; Stevens et al., 2016; Uddenberg, Akhter, Ramachandran, Sundström, & Carlsbecker, 2015; Wan et al., 2018; Zimin et al., 2014), has slowed down comparative genomic analyses in conifers with angiosperms (Chen et al., 2018).

Given this limited availability of whole genome reference sequences, the vast majority of current genomic analyses in conifers rely on *de novo* assembled reference transcriptomes obtained with next generation sequencing of RNA (RNA-Seq) (Baker et al., 2018; Z. Li et al., 2015; López de Heredia & Vázquez-Poletti, 2016). The use of these assembled references in conifer trees, therefore, has expanded to other applications in addition to the identification of differentially expressed genes (DEGs). These applications include comparative genomic analyses (Baker et al., 2018; De La Torre, Li, Van De Peer, & Ingvarsson, 2017; Wachowiak, Trivedi, Perry, & Cavers, 2015), and expression-QTL mapping (Verta, Landry, & Mackay, 2016). It also includes phylogenomic analyses (Li et al., 2017), SNP discovery and molecular marker development (Canales et al., 2014; Parchman, Geist, Grahnen, Benkman, & Buerkle, 2010), and gene identification for exome-capture bait development (Howe et al., 2013; Müller, Freund, Wildhagen, & Schmid, 2015). However, the generation of reliable and complete reference transcriptomes still faces several challenges. Because many non-model species are outbreeding, their genome displays high levels of heterozygosity that hampers *de novo* assembly algorithms and causes allelic redundancy (the presence of alleles of the same gene on different transcripts) and transcript fragmentation (splitting of portions of the same gene). Hence, these reference transcriptomes usually contain a larger number of contigs (transcripts) than the number of expressed genes (Gayral et al., 2013; Ono et al., 2015).

Several strategies have been employed to handle allelic redundancy and transcript fragmentation in *de novo* assembled transcriptomes(Davidson, Hawkins, & Oshlack, 2017; Davidson & Oshlack, 2014; L. Fu, Niu, Zhu, Wu, & Li, 2012). These approaches include scaffolding translation mapping (STM) during the assembly (Surget-Groba & Montoya-Burgos, 2010), post-scaffolding methods after assembly (TransPs) (Liu, Adelman, Myles, & Zhang, 2014), and Orthology Guided Assembly (OGA) (Ruttink et al., 2013). It includes as well the identification of orthologous contigs using partial or complete genome sequences (Armero, Baudouin, Bocs, & This, 2017; Bao, Jiang, & Girke, 2013), and phylogeny-informed identification of orthologous sequences (Medlar, Laakso, Miraldo, & Löytynoja, 2016). Although some of these strategies have been successfully applied to some crop species (e.g., Stočes et al., 2016), these methods rely on the quality of assembled and annotated genomic resources from closely related species to cluster allelic sequences and scaffold fragmented transcripts. In conifers, these methods have not been applied so far and allelic redundancy of assembled transcriptomes in this group is commonly reduced with CD-HIT clustering and the selection of the longest representative, or with CAP3 clustering (Li et al., 2017; Suren et al., 2016). Thus, the levels of allelic redundancy of previous published transcriptomes in conifers has not yet been determined, it is unclear to what extent allelic redundancy, and transcript fragmentation influences downstream applications. Considering the diverse range of applications of *de novo* reference transcriptomes in conifer trees (Canales et al., 2014; Celedon et al., 2017; De La Torre, Lin, Van De Peer, & Ingvarsson, 2015; Hu et al., 2016; Pinosio et al., 2014; Porth, White, Jaquish, & Ritland, 2018; Raherison et al., 2012), there is currently a need for additional strategies to assess the suitability of a transcriptome assembly for some of these diverse applications.

Paralog sequence collapse (PSC), the over-clustering of highly similar sequences (meant to collapse allelic sequences belonging to the same gene) leads to another type of problems in assembly and its downstream applications. These occur in the original assembly when different paralogs are assembled to create for instance mosaics. Allelic redundancy reduction will increase the frequency of PSC. However, most plant genomes contain a number of closely related gene copies (paralogs) due to whole genome or single gene duplications, making PSC and consequential paralog mapping (the mapping of reads originating from different paralogs on the same location of the reference) a common problem across species. In evolutionary and population genetic analyses, it is desirable to keep even highly similar gene copies separate, since after the duplication event they obtained distinct genomic locations, gene genealogies and evolutionary histories. Collapsing of paralogous gene copies will lead to paralog mapping and false variant calls. This redundancy is originated from allelic sequences that are assembled together or the scaffold of neighboring contigs derived from a fragmented assembly. These artifacts are usually addressed during redundancy reduction. However, if paralogous sequences are collapsed into a single representative sequence in the reference during redundancy reduction, reads originating from those different gene copies will map together and variation between two paralogs will appear as polymorphism. One solution for this problem is the identification and exclusion of paralogs after the SNP calling with model based approaches or by identification of excess heterozygosity (Gayral et al., 2013; McKinney, Waples, Seeb, & Seeb, 2017), which may lead to considerable loss of data in cases when PSC is common.

Here, we take advantage of the availability of different tissue ploidy levels in conifers to determine the effects of allelic redundancy and PSC on downstream analyses. Using the single-seed haploid megagametophyte tissue in *Pinus sylvestris* L., we first tested three strategies to generate *de novo* assembled transcriptome references and second, used a novel approach to estimate levels of paralog mapping (Figure 1). PSC and the resulting paralog read mismapping in the context of this study was studied by utilizing the expected lack of heterozygosity in samples obtained from haploid megagametophyte tissues. The amount of observed haploid heterozygosity (H_o_) in various assembled transcriptomes in relation to observed and expected heterozygosity (H_E_) in diploid tissues was then used to assess the levels of PSC and its effects on estimates of genetic diversity in each assembly. We further discuss the effect on each strategy used to generate a reference transcriptome and how the levels of PSC can influence downstream applications. Particularly, we focused on the effect of PSC of reference transcriptomes for studies aiming for: 1) marker development (especially for probe design and the availability of a non-redundant reference sequence for downstream mapping analyses), and 2) use of the assembled transcriptome as a reference for population genetic analyses and genetic diversity estimates.

**Figure 1.**
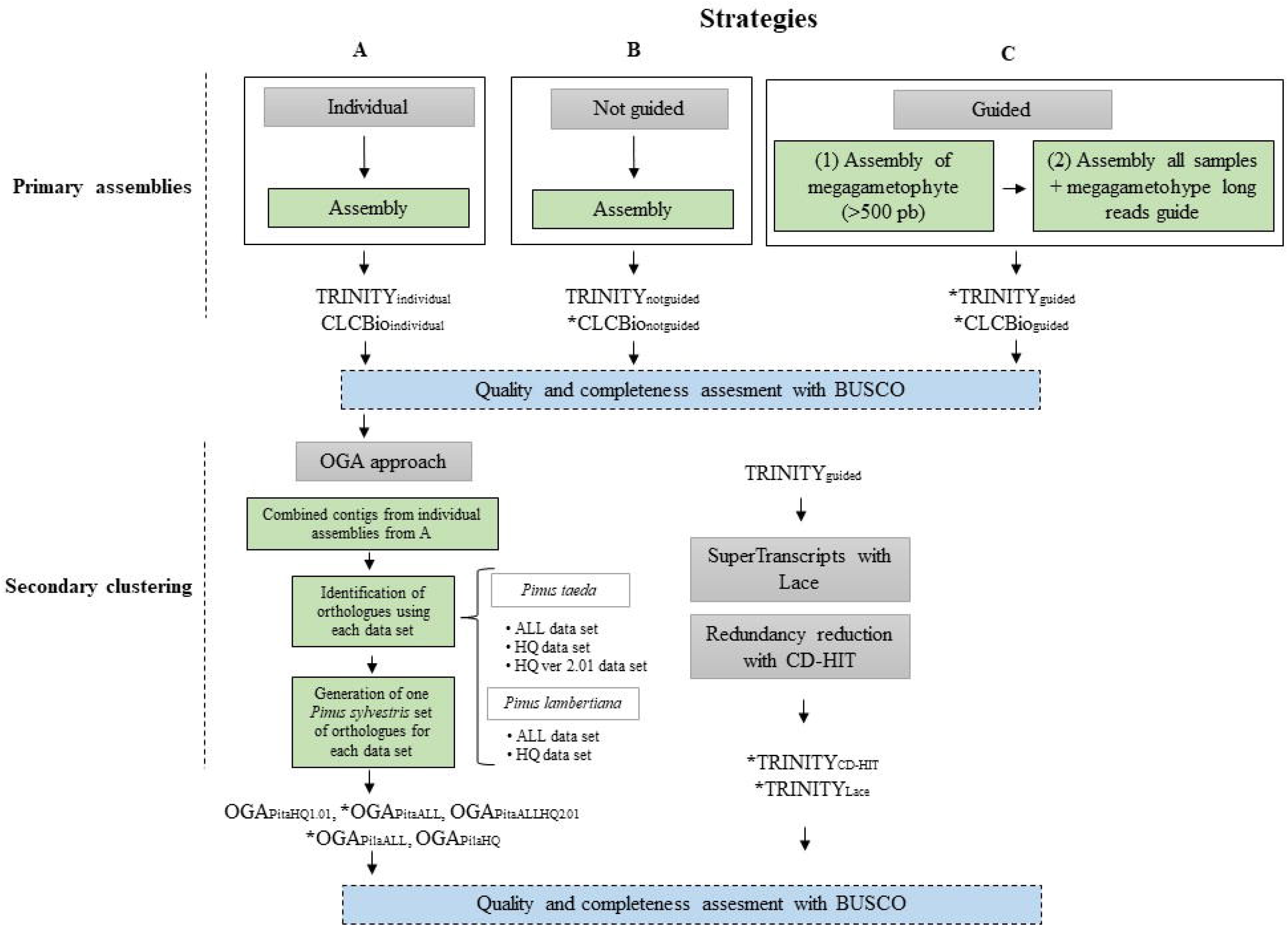
Strategies used to generate and evaluate the reference transcriptome for *P. sylvestris.* A) Individual assemblies, B) combined assembly of all reads per sample, and C) assembly of all megagametophyte (ME) per sample; retaining only > 500 bp transcripts (1) and then all different tissues per sample combined using the ME assembly as guidance sequences during the *de novo* assembly (2). Trinity and CLCbio Workbench assemblers were used on all three strategies. The secondary clustering consisted of the Orthology Guided Approach (OGA), construction of Lace SuperTranscripts followed by CD-HIT reduction of allelic redundancy. Assemblies marked with an * were assessed for levels of paralog mapping

## 2. MATERIALS AND METHODS

### 2.1 Plant material

We collected five tissues (needle, phloem, vegetative bud, embryo and megagametophyte) from six non-related individuals of *P. sylvestris* growing in a forest study site located in the Punkaharju, Southern Finland on May 26th -27th, 2016 (Table S1). At the time of collection, male strobili were shedding pollen and the vegetative growth for the year had already started. The sampled trees grow in two close-by naturally regenerated locations (Mäkrä and Ranta-Halola, Finland). Needles (NE), phloem (PH) and vegetative buds (VB) were dissected in the field and stored immediately in RNAlater (Thermo Fisher Scientific). Samples were transported on dry ice and kept in -80°C or -20°C (samples in RNAlater) until RNA extraction. Megagametophytes (ME) and embryos (EM) were sampled from a single germinating seed of the abovementioned mother trees. Before germination, seeds were stored in the dark at 4°C. Germination was initiated by keeping the seeds in moist paper, under constant light (300 umol/m^2^/s) and in 23°C degrees for 48 hours. Each seed was carefully dissected by removing first the seed coat, the nucellar cap and layers, and taking care of separating the diploid embryo and haploid megagametophyte tissues. From each seed, megagametophyte tissue and embryo were collected, and rinsed with 70% ethanol during the dissection.

### 2.2 RNA isolation, library preparation and sequencing

mRNA was directly extracted from the embryo (EM) and megagametophyte (ME) using Dynabeads mRNA Direct Micro Kit according to manufacturer’s instructions, except for a minor modification (using 200 μl of lysis buffer). Total RNA was extracted from needles (NE), vegetative buds (VB), and phloem (PH) with the Spectrum Plant Total RNA Kit (Protocol B, Sigma). After total RNA extraction, mRNA was captured using NEBNext^®^ Poly(A) mRNA Magnetic Isolation Module (New England Biolabs Inc.). mRNA extractions were treated with Turbo DNA-free Kit (Thermo Fisher Scientific). RNA concentration was quantified using Qubit 2.0 (Invitrogen) and Qubit^®^ RNA HS Assay Kit (Thermo Fisher Scientific). All libraries (6 trees x 5 tissues) were prepared using the NEBNext Ultra Directional RNA Library Prep Kit for Illumina (New England Biolabs Inc.) with a fragmentation time of 5-12 min. An insert size selection of 300 bp was targeted using a concentration of 40-45 μl per 20 μl AMPure XP (Agencourt) and between 12-15 cycles of PCR during library preparation. Libraries were indexed using NEBNext Multiplex Oligos for Illumina, Single Index Set 1. RNA library concentrations were quantified using NEBNext^®^ Library Quant Kit for Illumina and LightCycler 480 (Roche). Fragment size distributions of mRNA, total RNA and libraries were verified with 2100 Bioanalyzer RNA 6000 Pico Kit and DNA 1000 kits (Agilent). 6-12 libraries were pooled in 5 runs of an Illumina NextSeq500 instrument and sequenced with Mid-Output Kit (Illumina) in the Biocenter Oulu Sequencing Center.

### 2.3 Strategies to generate *de novo* reference transcriptomes: primary assemblies

Raw read quality was analyzed with FastQC (Andrews, 2010), and adapters and low quality reads were removed with Trimmomatic 0.33 (Bolger, Lohse, & Usadel, 2014) using the following parameters: “TruSeq3-PE-2.fa:2:30:10:1:true LEADING:3 TRAILING:3 SLIDINGWINDOW:10:20” We used three main strategies to perform primary reference assembly for *P. sylvestris.* The first primary assembly is based on individual assemblies, using all reads per sample per genotype using CLC Genomic Workbench (www.clcbio.com) ver. 10.1 with default settings or Trinity ver.2.4.0 with default settings (with the - group-pairs-distance 1000) (Haas et al., 2013). This resulted in 30 separate assemblies per algorithm, here called TRINITY_individual_ and CLCbio_individual_) (strategy A in Figure 1). The second strategy first combined all reads from all samples (700 × 10^6^ reads), normalized the reads (default 50 coverage) and performed *de novo* assembly using CLCbio and TRINITY with “not guided” mode (strategy B in Figure 1, further referred to as CLCbio_notguided_ and TRINITY_notguided_). The third strategy first performed individual *de novo* assemblies, using CLCbio and TRINITY, for each of the six haploid megagametophyte tissues where only a single allele per gene is expected. We retained only transcripts > 500 bp, thus generating pseudo long reads for subsequent guided assembly. These >500 bp transcripts from the single-seed megagametophyte tissue are used for resolving isoforms, and improving assembly of complex transcripts, but they are not incorporated into the final assembly (https://github.com/trinityrnaseq/trinityrnaseq/wiki) (strategy C in Figure 1, referred to as CLCbio_guided_ and TRTNITY_guided_). Assemblies obtained from both programs using the three strategies (A, B and C) were deposited in the NCBI TSA repository, under BioProject XXX.

### 2.4 Quality assessment and completeness of primary assemblies

The assemblies obtained from the three above described strategies were evaluated in terms of number of contigs, N50, average contig size and number of predicted ORFs. These statistics were obtained from TrinityStats, rnaQuast (Bushmanova, Antipov, Lapidus, Suvorov, & Prjibelski, 2016), and TransRate (Smith-Unna, Boursnell, Patro, Hibberd, & Kelly, 2016). We further calculated the N50 considering only the transcripts with an expression value that represented 90% of the total expression data (E90N50). This was done for the Trinity assembled transcriptomes only (TRINITY_guided_ and TRINITY_notguided_) based on their levels of expression. First, the individual reads were separately mapped to each reference using eXpress (Roberts & Pachter, 2013), then the matrix of counts was generated using the scripts of the Trinity RNAseq pipeline (Supplementary Data S1). Additionally, a metric of gene completeness of all these assemblies was determined with BUSCO ver 2.0 using the embryophyta_odb9 database(Simão, Waterhouse, Ioannidis, Kriventseva, & Zdobnov, 2015) (Figure 1).

### 2.5 Secondary clustering: the Orthology Guided Assembly (OGA) approach

After the assessment of the individual transcriptomes obtained in strategy A, we selected the reference from one assembler (CLCBio_individual_) and combined all the contigs generated from the individual assemblies. This combined set of *P. sylvestris* contigs was used for secondary assembly using OGA (Ruttink et al., 2013) with previously published proteomes from *Pinus taeda* and *Pinus lambertiana* (Figure 1) to guide the assembly. We either used 1) all annotated proteins (ALL) from *P. taeda* v1.0; or 2) all annotated proteins (ALL) from *P. taeda* v2.01 (Neale et al., 2014); or 3) all annotated proteins from *P. lambertiana* v1.0 (Gonzales-Ibeas, Martinez-Garcia, Famula, & Delfino-Mix, 2016; Stevens et al., 2016); or 4) only the high quality curated proteins (HQ) from *P. taeda;* or 5) from *P. lambertiana* (Table 1). Briefly, OGA first uses sequence similarity (tBLASTn) to the proteomes of the reference species to select allelic and fragmented contigs from all genotypes (assembled individually) per reference protein, then applies CAP3 clustering on a gene-by-gene basis (Ruttink et al., 2013), and finally selects the most likely orthologous CAP3 contigs per protein of the reference species. With this procedure it is possible to resolve transcript fragmentation and allelic redundancy across the individual assemblies, while generating a transcriptome reference sequence for *P. sylvestris* representing orthologs of a closely related species. The set of *P. sylvestris* orthologs identified with the OGA approach with all five data sets are deposited in the NCBI TSA repository, under BioProject XXX.

**Table 1.**
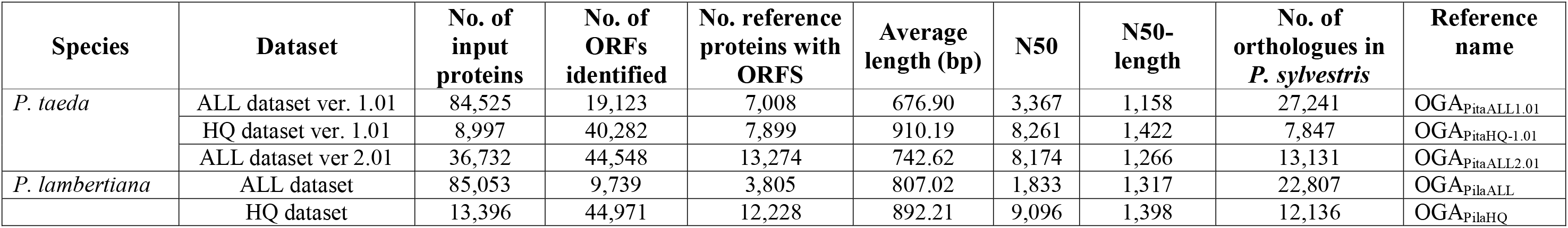
Proteomes used as reference for Orthology Guided Assembly (OGA) of *P. sylvestris* transcriptomes. Number of input proteins used per reference species and the number of orthologs identified in *P. sylvestris* with the OGA approach.

### 2.6 Secondary clustering: Construction of SuperTranscripts with Lace and CD-HIT

The different isoforms assembled for each gene in the TRINITY_guided_ reference were clustered into SuperTranscripts (representation of all isoforms by a single non-redundant contig) using the Lace software version 1.00 (Davidson et al., 2017). Briefly, we first divided the data into separate files with Lace based on cluster information from the TRINITY assembly and then generated multiple alignments for each cluster using BLAT v.35 (Kent, 2002). Based on the multiple alignment of the isoforms, we applied a graph-based algorithm to generate a single sequence (SuperTranscript) containing combined information from all isoforms (Davidson et al., 2017). Hence, each SuperTranscript ideally represents a single genomic locus. To avoid spurious isoforms with very low support, we excluded short (≤ 300 bp) and low expressed (total effective count ≤ 10 read count) isoforms prior to running Lace. In cases where all isoforms per cluster were filtered out after these exclusion criteria, the longest isoform was kept in the reference to avoid total exclusion of this sequence from the resulting reference transcriptome. We further identified SuperTranscripts that appeared to contain similar exonic sequences merged consecutively, as these kinds of “mosaics” could arise due to allelic variation and PSC. These were identified by self-blasting the SuperTranscripts. All SuperTranscripts that had blast hits within itself (other than the obvious 100% self-match) were identified as potential mosaics

Additionally, in order to inspect the effect on commonly applied clustering procedures that are used to decrease the allelic redundancy of assembled transcriptomes, the Trinity assembly selected (TRINITY_guided_) was clustered using CD-HIT-EST version 4.7 ((L. Fu et al., 2012; Weizhong Li & Godzik, 2006) with sequence identity cut-off (-c) 0.95, a commonly used threshold (Hodgins, Yeaman, Nurkowski, Rieseberg, & Aitken, 2016; Li et al., 2017; Wachowiak et al., 2015). The assemblies of *P. sylvestris* obtained using these two additional secondary clustering strategies are deposited in the NCBI TSA repository, under BioProject XXX.

### 2.7. Quality assessment and completeness of secondary clustering references

Finally, we evaluated the reference transcriptome assemblies obtained from the secondary clustering with Transrate (Smith-Unna et al., 2016) and estimated their completeness with BUSCO using the embryophyta_odb9 database (Simão et al., 2015) (Figure 1). After this assessment, we selected three of the references obtained with the secondary clustering (Figure 1, references marked with an *), which were used as reference transcriptomes to map the haploid (megagametophyte) and diploid tissues for assessment of the levels of PSC.

### 2.8 Assessment of the levels of PSC in references transcriptomes using haploid and diploid tissues

In order to evaluate the level of PSC on the obtained reference transcriptomes, we utilized the known ploidy level of two tissue types, the haploid (megagametophyte) and diploid tissues. In total, we employed six independent individuals, representing three different genotypes per individual: the vegetative tissues needle, phloem and vegetative bud (pooled reads of NE, PH, VB per individual) representing the diploid maternal genotype. The megagametophyte (ME) material containing a haploid maternal genome while the embryo (EM), being diploid but representing the next generation. One of the haploid genomes of the embryo is the same as the seed megagametophyte genome. The other haploid genome comes from an unknown pollinating father. Individual reads from ME, EM and NE+PH+VB of these six individuals were mapped against the seven selected reference transcriptomes obtained, four from the primary assemblies and another three obtained with the secondary clustering (Figure 1, assemblies with an *). Mapping of the reads was performed with STAR aligner (Dobin et al., 2013) given its suitability to allow spliced mapping of reads originating from RNA sequencing, e.g. super transcriptome reference combining multiple isoforms (Dobin et al., 2013). We used default settings, with the exception of allowing reads to map to only one locus in the reference (–outFilterMultimapNmax 1). Otherwise, the read was considered unmapped. We also modified the default setting for filtering alignments with a mismatch of 0.025 (–outFilterMismatchNoverLmax 0.025). We used the two-pass mapping strategy (– twopassMode Basic) where STAR first performs the first pass mapping, then extracts junctions, inserts them into the genome index, and finally uses this information during the remapping of all reads in the second mapping step. Duplicates were removed with SAMtools (H. Li et al., 2009) and read groups added with the picard tool AddOrReplaceReadGroups. We generated three vcf files per reference transcriptome according to the ploidy of the tissues: one vcf for the ME (haploid), one for the EM (diploid), and one combining NE+PH+VB (diploid). Monomorphic and polymorphic sites were called with FreeBayes (Garrison & Marth, 2012) using default parameters, with exceptions of using a mutation rate (-T) of 0.005, excluding indels (-i), ignoring complex events (-u), and allowing no MNPs (multi-nucleotide polymorphisms) (-X). Each vcf file with both monomorphic and polymorphic sites was filtered with vcftools (Danecek et al., 2011) with a minimum depth per sample set to 10, and maximum amount of missing data 0.5 per site. Further filters were applied to polymorphic sites keeping only bi-allelic SNP sites with a quality > 20. Number of heterozygous and homozygous variant calls were determined with vcftools –hardy option. Number of callable sites, sites with sufficient depth and amount of missing data per variant was determined for each transcript to allow comparable, per bp level nucleotide diversity estimates. Both observed and expected heterozygosity (π) (Tajima, 1989) per nucleotide was calculated for each transcript individually. Given that the expected value of π is equal to θ (=4N_e_μ), we contrasted our observations to earlier, independent θ estimates for *P. sylvestris* (Grivet et al., 2017; Kujala & Savolainen, 2012; Pyhäjärvi et al., 2007; Pyhäjärvi, Kujala, & Savolainen, 2011). Heterozygous calls from reads originating from haploid tissues indicate paralog mapping or sequencing/genotype calling errors and we used this to compare the level of PSC among the reference transcriptomes. The workflow with all the commands used is deposited in https://github.com/DI-Ojeda/Pin_syl_transcriptome.

### 2.9 Functional annotation and identification of contaminants on the assembled transcriptomes

The assembled transcriptomes selected as references for differential gene expression (TRINITY_guided_, see criteria later) was annotated using Trinotate, a pipeline for functional annotation of transcriptomes (Bolger, Arsova, & Usadel, 2017). First, similarities to known proteins were detected by a BLASTX search (Camacho et al., 2009) (e ≤ 1e–5) against two comprehensive protein databases: Swiss-Prot (Boeckmann et al., 2005) and UniRef90 (The UniProt Consortium, 2015) obtained from UniProt (available on Mar 8, 2018). Coding regions within transcripts were predicted using TransDecoder (http://transdecoder. github.io). The protein products identified from TransDecoder were searched for sequence similarities against the Swiss-Prot and UniRef90 protein database and for conserved protein domains using Hmmer (http://hmmer.org/) and PFam (Finn et al., 2014). All results were parsed by the Trinotate pipeline (https ://trinotate. github.io), stored in a SQLite relational database, and reported as a tab-delimited transcript annotation file.

We also examined whether possible contaminants were present in the final assemblies. In order to identify contaminant sequences (not belonging to *P. sylvestris*), we performed a BLASTx search of the assembled reference (TRINITY_guided_) against Swiss-Prot limited to sequences classified as bacteria, viruses, metazoan, alveolata and/or fungi. We identified transcripts that had a BLASTx hit e ≤ 1e–5 and sequence similarity of at least 65%. Transcripts potentially originating from organelle genomes were identified by BLAST against P. taeda and P. lambertiana mitochondrial genomes and *P. sylvestris* and *P. mugo* chloroplast genomes (NCBI GenBank IDs JN854158.1 and KX833097.1) with e-value cutoff 5e-2, identity cutoff 80 % and word size = 60. BLAST search of the transcripts against the *P. taeda* (v.2.01) (Zimin et al., 2014) and *P. lambertiana* (v.1.0,) (Stevens et al., 2016) reference genome sequences was used as an additional method to identify single-copy and multi-copy transcripts. If any region of a given transcript had multiple BLAST hits with more than 85% identity, the transcript was assigned as multi-copy. We excluded 10 bp from each edge of the alignments to avoid random alignment in the edges. In addition, we required that 50% of the transcript length had sequence similarity in the corresponding reference.

### 2.10 Completeness of assembled transcriptomes and comparisons with published transcriptomes of *P. sylvestris*

In order to determine the completeness of the transcriptome references obtained for this species, we used the core set of genes (embryophyta_odb9) dataset in BUSCO (Simão et al., 2015). Although this core set is only based on angiosperm taxa, it provides an estimate for the completeness of a core set of genes, also for gymnosperms. We evaluated three of the references obtained here (TRINITY_guided_, TRINITY_CD-HIT_, and TRINITY_Lace_), and compared them to previously published assemblies for *P. sylvestris.* Transcriptomes for this species have been assembled from heartwood of wounded and unwounded seedlings (Godfrey, 2012), needles of two-year-old seedlings collected from five individuals (genotypes) (Wachowiak et al., 2015), embryos and megagametophytes at different developmental stages from a single individual (Merino et al., 2016). References are also available from wood cores collected from four individual mature trees (35 to 46-years-old) (Lim, 2017; Lim et al., 2016) and pollen (Höllbacher, Schmitt, Hofer, Ferreira, & Lackner, 2017).

## 3. RESULTS

### 3.1 Primary assemblies

Individual assemblies (Figure 1, strategy A) obtained with the CLCbio Genomic Workbench (CLCbio_individual_) contained overall less contigs (average 84,181) than TRINITY_individual_ assemblies (average 149,943) (Table S2) and displayed slightly (on average 5%) lower BUSCO completeness scores (Figures S1 and S2). We obtained similar results in terms of average contig size, longest assembled contig, and N50 among the four assemblies obtained with the pooled assembly strategies (Figure 1, strategies B and C) with Trinity and CLCbio (Table S3). A higher number of contigs was obtained with Trinity regardless if strategy B or C was used, and this was also reflected in the number of predicted ORFs. Using the megagametophyte > 500 bp contigs (Figure 1, strategy C) as a guide during assembly increased the total number of contigs obtained in the second assembly on both algorithms. A large proportion of transcripts assembled with Trinity have low expression values. For instance, we obtained 52,208 genes with a >10 TPM level of expression for the TRINITY_guided_ assembly (Figure S4). We obtained a slightly larger number of genes with the TRINITY_notguided_ strategy (64,986 genes, E90N50 = 2,158, the N50 value obtained with only a level of expression of 90%) than with the TRINITY_guided_ strategy (50,760 genes, E90N50 = 2,388) when a cut-off expression value of >90% was used (Figure S5). BUSCO completeness scores were also higher on these latter assemblies compared with strategy B. The highest BUSCO completeness scores and the lowest number of fragmented transcripts was obtained with the TRINITY_guided_ assembly (Figure S6A).

### 3.2 Secondary clustering: Orthology Guided Assembly (OGA)

We used the individual assemblies obtained with the CLCbio (CLCbio_individual_), which contained overall considerably lower complexity (measured by the number of unique contigs per sample, as function of the total number of reads available) (Figure S3). Given its reduced size compared to TRINITY assemblies, the identification of orthologs with the proteomes of *P. lambertiana* and *P. taeda* required less computational time. We combined the individual transcripts for each of the 30 samples resulting in a total of 2,495,509 transcripts (average length = 642.76 bp, N50-length = 930 bp) and used them as an input for the identification of orthologs with the OGA approach. Overall, we obtained a higher number of orthologs of *P. sylvestris* using the ALL proteome datasets for both reference species, with higher numbers with *P. taeda* (27,241 orthologs) than *P. lambertiana* (22,807 orthologs) (Table 1 and S5). This was also reflected in the BUSCO completeness scores, where a more complete core gene set was recovered with the ALL proteome datasets for *P. taeda* (42.6%) than for *P. lambertiana* (42.0%) (Table S4 and Figure S6B). The ability to recover a full-length *P. sylvestris* transcript per gene with the OGA approach was determined by plotting the distribution of the fraction of the assembled transcript length versus the respective *P. taeda* or *P. lambertiana* reference sequence length. We found that the majority of the transcripts obtained using the five reference proteomes (Table 1) were fragmented (broken or partially assembled transcripts) (Figure S7). For instance, using the latest version of the *P. taeda* protein set (Table 1, ALL dataset ver. 2.01) resulted in the recovery of 13,131 *P. sylvestris* transcripts (out of the 36,732 *P. taeda* reference proteins), but only one-third of the *P. sylvestris* transcripts encodes a full-length protein with a similar length as the *P. taeda* reference protein (ratio close to 1) (Figure S7).

### 3.3 Secondary clustering: Reduction of allelic redundancy with Lace and CD-HIT

After the previous assessments, the TRINITY_guided_ reference was further analyzed to reduce its redundancy with the construction of SuperTranscrips (Lace) and with CD-HIT. From the original set of 1,307,499 contigs in the TRINITY_guided_ assembly, a total of 787,820 SuperTranscrips were obtained with Lace, of which 71,344 (9 %) included multiple isoforms. Out of the multi-isoform clusters, most had a few (<5) isoforms (Table S5). With CD-HIT, we further reduced the number of transcripts by 9%, resulting in a set of 717,762 transcripts.

### 3.4 Assessment of levels of paralog sequence collapse (PSC)

To evaluate the level of paralog read mapping due to PSC on the reference transcriptomes obtained, patterns of heterozygosity among haploid and diploid tissues were compared on the seven selected reference transcriptomes (Figure 1). The summary of SNP and genotype calling results is presented in Table 2. It is noteworthy that both the size of the transcriptome and the number of callable sites (monomorphic and polymorphic sites passing the filters) are very different among assemblies, the latter varying from 6.7 × 10^6^ callable sites in OGA_pilaALL_ to 102 × 10^6^ callable sites in the TRINITY_CD-HIT_. Larger references resulted in lower average depth and vice versa, because smaller assemblies force the same amount of reads to map to smaller number of locations. In addition, the number of callable sites will act as a denominator in per nucleotide estimates, thus reducing the apparent level of diversity in the reference transcriptomes with more callable sites. Therefore, for evaluating the behaviour of different assemblers, we also considered the haploid/diploid ratio of H_o_ in addition to the absolute number of variants per nucleotide, as the ratio is less affected by the assembly size.

**Table 2.**
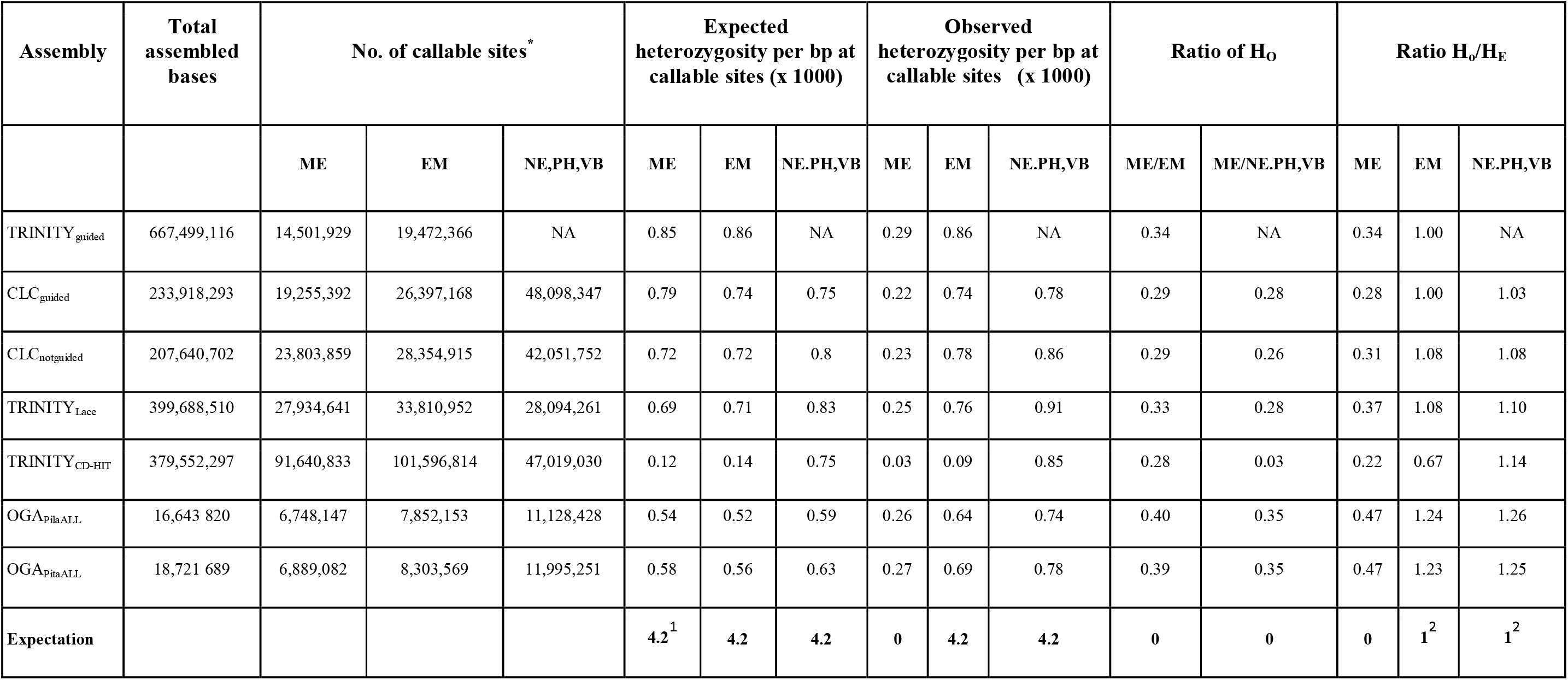
Summary of the assessment of PSC levels utilizing observed heterozygosity patterns on the seven reference transcriptomes selected. NA = not analyzed. *Only including genes where there are >100 callable sites. ^1^Rough estimate for total sites from Grivet *et al.*, 2017, assuming that one-third of coding sites are nonsynonymous and two-third are synonymous. ^2^Assuming Hardy-Weinberg equilibrium.

As expected, observed heterozygosity per nucleotide was lower in megagametophytes than in diploid tissues on all the assemblies assessed. However, some observed heterozygosity in haploid tissue in all assemblies suggest that all the assessed transcriptomes still result in considerable amount of false heterozygous calls, either originating from paralog read mapping or from errors during mapping and variant calling (Table 2). We found that the relative amount of false heterozygous calls (ratio of H_o_) was lowest in TRINITY_CD-HIT_ (TRINITY_guided_ assembly, followed by creation of SuperTranscripts and clustering redundant contigs with CD-HIT, Table 2). In contrast, haploid vs. diploid observed H_O_ ratio was the highest among all assemblies in OGA assemblies. Also the H_o_/H_E_ ratio was considerably higher than one for both OGA assemblies in diploid tissues suggesting excess heterozygote calls likely due to PSC.

As a further validation of our methods of identifying levels of paralog read mapping based on haploid heterozygosity, we compared the heterozygosity level in single and multi-copy genes identified in the TRINITY_Lace_ (Figure 1). We expected the paralog mapping and hence haploid observed heterozygosity to be higher in multi-copy genes that are prone to PSC. In diploid genotypes, multi-copy genes had slightly higher heterozygosity than single copy genes. In haploid genotypes, the majority (93%) of genes with H_o_ > 0 were found in multicopy genes (Figure S8). On the other hand, 30% of the multi-copy genes, but only 14% of the single-copy genes (where we obtained sufficient data for SNP calling) have H_o_ > 0. Thus, the two independent methods yield consistent but not completely overlapping methods to identify loci prone to PSC.

### 3.5 BUSCO completeness and functional annotation of the assembled transcriptome

Our final assemblies ranged between 61.0% and 82.7% completeness, with the highest level recovered in the TRINITY_guided_ assembly (Figure 2). In contrast to the previously published studies that assembled reference transcriptomes for *P. sylvestris* used only one or two different tissues (Höllbacher et al., 2017; Lim, 2017; Lim et al., 2016; Merino et al., 2016; Wachowiak et al., 2015), our assembly is based on five different tissues (needle, phloem, vegetative bud, embryo and megagametophyte) from six individuals. The number of full-length genes captured in the TRINITY_guided_ assembly is also reflected in the number of ORFs predicted with TransRate, which was larger than the previously available reference for *P. sylvestris* (Table S6). The TRINITY_guided_ reference therefore represents the most complete available resource for this species to date. Using the Trinotate pipeline we annotated the 1.3 million transcripts of this assembly and found a large number (50,760) of likely full-length transcripts enriched in the E90 set (Figure S5a) and at least 64,260 complete ORFs predicted with Transdecoder. In addition, Trinotate found 24,780 genes matching Pfam Tracheophyta protein hit (Supplementary Data S1).

**Figure 2.**
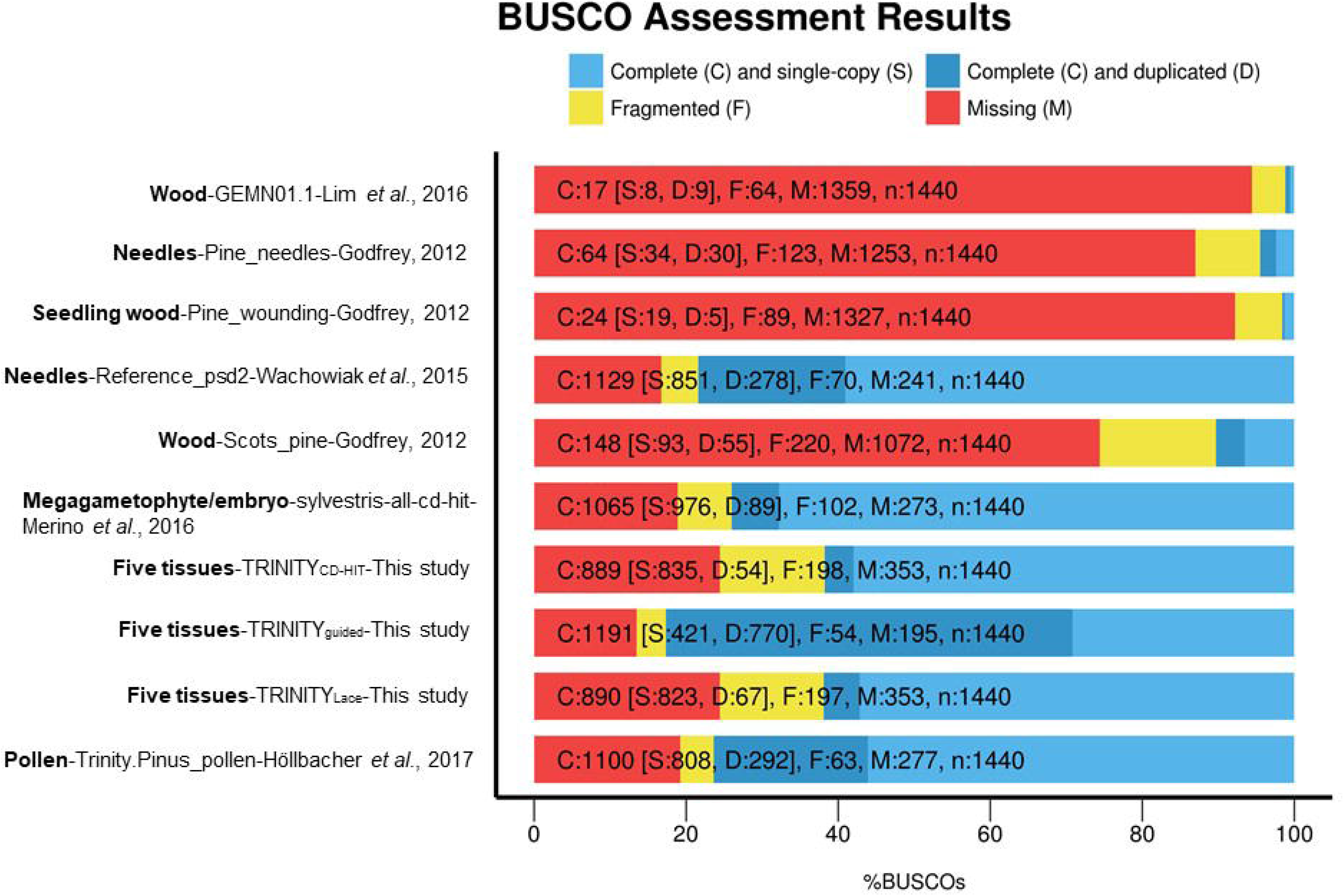
Percentage of completeness on the core set of genes in BUSCO of the assemblies obtained in this study in comparison with published transcriptomes.

## 4. DISCUSSION

Considering the large and complex genomes of conifers, reference transcriptomes are increasingly being used as a reference resource in a variety of applications (Müller et al., 2015; Suren et al., 2016; Wachowiak et al., 2015). Particularly in Pinaceae, where the vast majority of transcriptomes have been generated in gymnosperms (López de Heredia & Vázquez-Poletti, 2016), the number of assembled contigs (transcripts) is always larger (Table S6) than the actual number of genes estimated based on their genome sequence (Gonzales-Ibeas et al., 2016; Neale et al., 2014; Nystedt et al., 2013).

One of the main limitations and potential source of error is the level PSC in the reference transcriptome, whereby similar, but paralogous gene sequences are erroneously collapsed into a single reference sequence. One particular characteristic of conifers (and other gymnosperms) is the presence of easily accessible megagametophyte tissue, and their amenity to extract haploid genome information from it. We used the variation of ploidy in *P. sylvestris* tissues to assess their utility during the *de novo* assembly and employ a novel strategy to estimate levels of PSC on the assembled transcriptomes. The same approach could be utilized in any species with haploid material available (e.g., social insects, haplontic plants and fungi).

### 4.1 Improving *de novo* assembly transcriptomes with the haploid megagametophyte tissue

The most common strategy applied in *de novo* transcriptome assembly involves combining read data from several genotypes (individuals), developmental stages, and tissues of the target species. This is justified as a mean to capture most of the genes expressed under a variety of conditions and individuals. However, this also causes high levels of allelic redundancy and transcript fragmentation due to the high levels of heterozygosity and the genetic diversity across several genotypes (Ruttink et al., 2013). In conifers, this is the strategy routinely employed to generate *de novo* reference transcriptomes (López de Heredia & Vázquez-Poletti, 2016), and it invariably leads to a large number of transcripts. Among the three strategies used here (Figure 1), combining all tissues and genotypes (strategy B) resulted in an assembly with the highest number of assembled contigs and predicted ORFs, but without necessarily being the most complete assembly (Table S3). On the other hand, the inclusion of the megagametophyte (haploid) during the *de novo* assembly as long read guidance (Figure 1C), increased the BUSCO completeness scores for both assemblers (Table 2). Contrary to *de novo* transcriptome hybrid assembly, where long reads (such as PacBio) are aligned to the short reads during the assembly (S. Fu et al., 2018), the long reads we used here from the megagametophyte (>500 bp) were not incorporated into the final assembly, and were only used to resolve isoforms during assembly. The main benefit obtained from incorporating haploid tissue reads during the assembly stage is the increase of transcriptome completeness. Without considerably affecting the number of transcripts or the ORFs predicted, regardless of the assembler used (Table S3), which were still considerably larger than the estimated number of genes in conifer genomes (Neale et al., 2017; Nystedt et al., 2013; Stevens et al., 2016; Zimin et al., 2014).

### 4.2 Levels of paralog sequence collapse in the assembled transcriptomes and the effect of redundancy reducing strategies

In conifers, CD-HIT clustering is the most common strategy used to reduce allelic redundancy, along with CAP3, with similar levels of effectiveness using either strategy (Yang & Smith, 2013). In contrast, generation of gene-based consensus transcript using isoform alignment (Lace) has been rarely applied in conifer trees (Ueno et al., 2018). The CD-HIT algorithm was originally designed to reduce large protein data sets into representative sequences based on homology (W. Li, Jaroszewski, & Godzik, 2001), not to reduce redundancy in transcriptomes. In our study, CD-HIT reduced the number of transcripts and predicted ORFs more than the generation of SuperTranscripts with Lace alone (Table S4). Both Lace and CD-HIT reduced the percentage of duplicated genes in the BUSCO set to about 5% of the genes (Figure 2). As a side effect, applying these clustering steps reduces the BUSCO completeness scores of the transcriptomes by nearly 20% in *P. sylvestris* (Table S4). In addition, Lace resulted in the inclusion of duplicated exons on the generated sequences, resulting in the generation of artificial mosaic transcripts. We identified a total of 44,535 mosaics on this assembly before applying the CD-HIT clustering. Contrary to CD-HIT and Lace, the OGA approach was less successful in the reduction of false heterozygous calls. For OGA assemblies, the observed heterozygosity was 25% higher than the expected heterozygosity, indicating a higher number of false heterozygous calls (Table 2). This was mainly due to the lack of completeness of the *P. taeda* and *P. lambertiana* reference genome annotations. As better and more complete reference genomes become available in conifers, this approach might be an effective strategy to reduce allelic redundancy, PSC and paralog read mapping.

### 4.3 Effect of allelic redundancy on the development of exome capture probes and strategies to mitigate its effect

Appropriate selection of target genes is a crucial step for an effective target exome capture experiment, as the presence of paralog sequences, misassembled regions, mosaics, cpDNA and mtDNA decrease the efficiency of the recovery of the captured regions. When the design is entirely based on an assembled transcriptome, the most important step is to select the most appropriate contigs for bait design. For those references obtained with programs that group assembled contigs by graph component (multiple isoforms) such as Trinity, a representative sequence must be selected for bait design. This is usually accomplished by selecting the isoform with a minimum length (Suren et al., 2016), or combining information from isoform size and their expression level (Yang & Smith, 2013). Based on our estimates of PSC, collapsing paralogs with Lace and CD-HIT after a haploid-guided assembly is a reasonable strategy to select suitable candidate genes. However, additional considerations should be added to further select the most suitable regions for bait design (Figure 3). Additional steps must include the removal of contaminant regions (Howe et al., 2013), genes from the organelle genomes (Syring et al., 2016), exclusion of potential mosaics produced by Lace, and the exclusion of transcripts with low level of expression. In our case, 0.4 % of the transcripts had a significant hit to chloroplast or mitochondrial Pinus genomes in the TRINITY_Lace_ assembly. Excluding genes with low levels of expression (cut-off of >10 TPM) reduced the number of genes (including their isoforms) to 108,860 after the TRINITY_guided_ assembly (Figure S5B). These low expressed genes are probably enriched with assembly artifacts (genes with no biological significance) and therefore not desirable in the bait design. For future reference and to help exome capture bait design of *P. sylvestris*, information on copy number, organelle genome match, isoform number, expression, putative contaminants and mosaics at gene level for Trinity_Lace_ gene level assembly is provided in Data S1.

**Figure 3.**
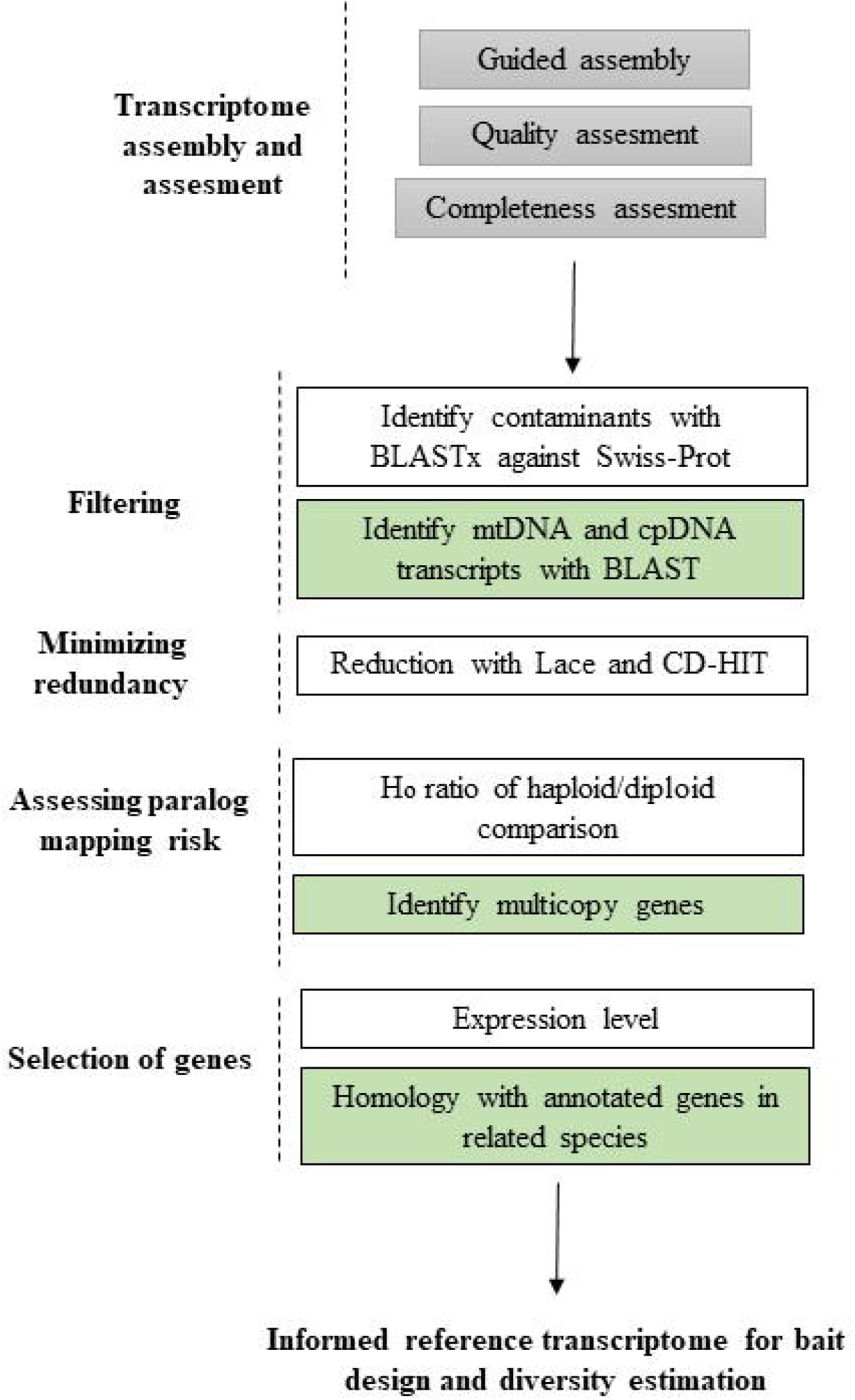
Strategies to employ the availability of haploid tissue in conifers for *de novo* transcriptome assembly and their application to assess the amount of allelic redundancy and paralog sequence collapse (PSC). Green boxes indicate additional steps recommended when additional genomic resources are available from a related species.

One aspect that has been rarely explored in the assembled transcriptomes of conifers is the proportion of chimaeras, which has been estimated between 4-9 % in the assembled transcriptomes evaluated so far (Ueno et al., 2018). There are currently available strategies to identify chimeras in assembled transcriptomes (e.g. Yang and Smith, 2013), but these rely on the quality of annotated proteomes from closely related taxa. Thus, how effective this strategy is in conifers remains to be determined. Finally, the intron-exon boundaries of the selected genes can be inferred using annotated genome references of related species, and excluded from the target regions of the bait design, as it has been shown to improve capture efficiency due to the hybridization on the probe-exon regions (Neves, Chamala, Davis, Barbazuk, & Kirst, 2013; Suren et al., 2016). However, the success of predicting these boundaries rely on the quality of available reference genomes from closely related taxa or low coverage shotgun sequencing of the target species. Depending on the availability of genomic resources from closely related taxa, we suggest several tailored steps that can be integrated to select suitable target genes for bait design (Figure 3).

### 4.4 Influence of transcriptome assembly strategies on estimates of genetic diversity

The use of transcriptomes as a reference for obtaining SNPs and estimating genetic diversity is becoming a common strategy for non-model species that lack a reference genome (Gayral et al., 2013; Romiguier et al., 2014; Xu et al., 2016; Yan et al., 2017). However, several factors can introduce errors, and careful assessment of the assembled transcriptome is required before employing this downstream application. These include the presence of alleles, isoforms, and paralogs on the assembled transcriptome contigs. For instance, allelic redundancy allows reads to map to their respective allele without mismatches, and because variant calling algorithms can report no polymorphisms, this will cause an underestimation of genetic diversity. On the other hand, too greedy clustering of transcripts will result in PSC, the collapse of paralogous sequences and the removal of some of these from the reference. As a result, reads derived from the genes that were removed from the reference may map onto the remaining sequence and variant calling algorithms will report nucleotide substitutions that differentiate the paralogous genes as SNPs (Gayral et al., 2013).

Overall, the H_e_ per nucleotide for the assembled transcriptomes we assessed was lower than expected based on previous studies (Grivet et al., 2017), ranging from 1.2 × 10^−4^ to 8.3 × 10^−4^ (Table 2), an order of magnitude lower than previously reported in *P. sylvestris* ((Pyhäjärvi et al., 2011). The minimum required read depth and missing data threshold we applied to our data set were relatively lenient, which has probably increased the number of callable sites relative to variant calls. Note that in this study the measures of diversity among haploid and diploid samples were used to evaluate different assemblies and were based on relative levels of heterozygosity. For precise estimates of genetic diversity, datasets with reads originating from DNA, deeper sequencing, more stringent filters and larger number of individuals should be used. We additionally suggest careful analysis of genetic diversity from multi-copy genes as they are especially prone to paralog mapping. Our estimation of heterozygosity on the haploid tissues were higher for multi-copy than single-copy genes (Figure S8), and the exclusion of multi-copy genes will decrease false SNPs in downstream analysis. This approach, however, comes with a caveat of possible bias resulting from ignoring fast evolving gene families from the analysis (Fig. 3).

## Supporting information

## Acknowledgments

We acknowledge the advice and help on bioinformatics from Lieven Sterck, John Liechty, Sam Yeaman, Henrik Hedberg, Tuomas Jormola, Jaakko Tyrmi, Witold Wachowiak and Nader Aryamanesh. Special thanks to Brian Haas for his helpful suggestions regarding the assemblies with Trinity. We thank Outi Savolainen and Komlan Avia for providing access to their experimental material. Soile Alatalo and Sami Saarenpää are acknowledged for skilled field and laboratory work. This work was funded by the Academy of Finland (287431 and 293819 to TP, GENOWOOD no. 307582), and the European Commission (EVOLTREE http://www.evoltree.eu/). We thank the CSC-IT Center for Science, Finland, for computational resources.

## Supplementary material

**Figure S1**. Percentage of completeness of the individual Trinity assemblies on the core set of genes in BUSCO.

**Figure S2**. Percentage of completeness of the individual CLCbio Genomics Workbench assemblies on the core set of genes in BUSCO.

**Figure S3**. Level of complexity (considered as the number of unique contigs per sample, as function of the total number of reads available) on the individual assemblies using Trinity and CLCbio Genomics Workbench. M= millions reads.

**Figure S4**. Distribution of the number of genes (A) and transcripts (B) based on their level of expression on the assembled transcriptome using Trinity and guided with the >500 bp contigs.

**Figure S5**. Graphical representation of the N50 values across the different levels of gene expression. The E90N50 values are shown for the Trinity assemblies using the guided strategy (A) and the not guided strategy (B).

**Figure S6**. Percentage of completeness on the core set of genes in BUSCO. A) The combined assemblies using CLCbio Genomics Workbench and Trinity with the guided and not guided strategies. B) The five assemblies obtained after the application of the OGA approach.

**Figure S7**. Success in the recovery of full-length orthologs (measured as the length ratio of *P. sylvestris* CDS length/reference CDS length) using the five reference proteomes of *Pinus lambertiana* (Pila) and *Pinus taeda* (Pita).

**Figure S8**. Distribution of H_o_ in the TRINITY_Lace_ reference between single-copy and multicopy (identified by BLAST against *P. taeda* genome) genes in A) haploid (megagametophyte), B) diploid embryo and C) diploid bud+needle+phloem tissues. D) is a detail of A).

**Table S1**. Sampling locations of the six trees used for RNA sequencing, *de novo* assembly and gene expression analysis.

**Table S2**. Statistics of the primary assemblies obtained with Trinity and with CLCbio Genomics Workbench.

**Table S3**. Statistics of the assemblies obtained with the combination of all reads using the strategies B and C with Trinity and CLCbio Genomics Workbench combining all reads.

**Table S4**. Statistics of the orthologs obtained with the OGA approach in *P. sylvestris* using the five data sets proteins from *Pinus lambertiana* and *Pinus taeda.*

**Table S5**. Number of SuperTranscripts classified based on isoform count.

**Table S6**. Statistics of previously published transcriptomes compared with transcriptome assemblies obtained in the current study.

**Data S1**. All 8 *Pinus sylvestris* transcriptomes, annotation file of the reference transcriptome obtained with the Trinity guided strategy (TRINITY_guided_), information on putative contaminants, organelle transcripts, mosaics, copy number and levels of expression of obtained with eXpress are available at:
https://pinus_sylvestris_transcriptome_public_data.object.pouta.csc.fi/Pinus_sylvestris_transcriptomes_Ojeda_2018.tar

