## Supplementary material for "Utilization of tissue ploidy level variation in *de novo* transcriptome assembly of *Pinus sylvestris*"

| **Tree ID** | **Latitude** | **Longitude** | **Location** |
| --- | --- | --- | --- |
| 251 | 61.655 | 29.280 | Ranta-Halola |
| 320 | 61.657 | 29.292 | Ranta-Halola |
| 397 | 61.838 | 29.396 | Mäkrä |
| 443 | 61.838 | 29.393 | Mäkrä |
| 463 | 61.837 | 29.392 | Mäkrä |
| 485 | 61.836 | 29.394 | Mäkrä |
