## Supplementary material for "Utilization of tissue ploidy level variation in *de novo* transcriptome assembly of *Pinus sylvestris*"

|  | **Combining all reads for the assembly (tissues x genotypes)** | | | |
| --- | --- | --- | --- | --- |
|  | **Strategy B, not guided** | | **Strategy C, guided** | |
| **Assembler** | **TRINITY** | **CLC Workbench** | **TRINITY** | **CLC Workbench** |
| **Reference name** | **TRINITY_notguided_** | **CLCbio_notguided_** | **TRINITY_guided_** | **CLCbio_guided_** |
| **Total assembled bases** | 681,954,381 | 207,640,702 | 667,499,116 | 233,918,293 |
| **Total number of contigs** | 1,288,196 | 517,592 | 1,307,499 | 549,976 |
| **Average of assembled contig** | 529.387 | - | 510.516 | 425.32 |
| **Longest transcript** | 17,466 | 16,627 | 18,287 | 11,207 |
| **GC content** | 40.39 | 41.00 | 40.38 | 41.00 |
| **N50** | 658 | 403 | 619 | 453 |
| **E90N50** | 2,158 | - | 2,388 | - |
| **BUSCO v 2.0.1 embryophyta_odb9** | C:71.9% [S:26.3%, D:45.6%] | C:54.5% [S:50.3%, D:4.2%] | C:82.7% [S:29.2%, D:53.5%] | C:60.8% [S:35.9%, D:24.9%] |
|  | F:8.7%, M:19.4%, n:1440 | F:16.8%, M:28.7%, n:1440 | F:3.8%, M:13.5%, n:1440 | F:14.8%, M:24.4%, n:1440 |
| **Transrate predicted ORFs** | 113,006 | 31,615 | 107,323 | 47,211 |
