## Supplementary material for "Utilization of tissue ploidy level variation in *de novo* transcriptome assembly of *Pinus sylvestris*"

| **Reference species** | *Pinus lambertiana* | | |
| --- | --- | --- | --- |
| **Reference name** | **OGA_PilaHQ_** | **OGA_PilaALL_** |  |
| **Total assembled bases** | 10,368,177 | 16,643,820 |  |
| **Total number of contigs** | 12,136 | 22,807 |  |
| **Average of assembled contig** | 841.03 | 708.72 |  |
| **Longest transcript** | 15,657 | 15,537 |  |
| **GC content** | 45.00 | 45.00 |  |
| **N50** | 1,401 | 1,293 |  |
| **BUSCO v 2.0.1 embryophyta_odb9** | C:31.7% [S:27.8%, D:3.9%] | C:42.0% [S:39.4%, D:2.6%] |  |
|  | F:7.1%, M:61.2%, n:1440 | F:3.6%, M:54.4%, n:1440 |  |
| **Transrate predicted ORFs** | 6,895 | 10,990 |  |
| **Reference species** | *Pinus taeda* | | |
| **Reference name** | **OGA_PitaHQ1.01_** | **OGA_PitaALL_** | **OGA_PitaHQ2.01_** |
| **Total assembled bases** | 6,723,369 | 18,721,689 | 11,663,547 |
| **Total number of contigs** | 7,847 | 27,241 | 13,131 |
| **Average of assembled contig** | 843.04 | 663.64 | 875.46 |
| **Longest transcript** | 7,359 | 11,019 | 11,676 |
| **GC content** | 45.00 | 45.00 | 44.00 |
| **N50** | 1,380 | 1,242 | 1,470 |
| **BUSCO v 2.0.1 embryophyta_odb9** | C:18.3% [S:16.2%, D:2.1%] | C:42.6% [S:39.0%, D:3.6%] | C:17.2% [S:15.3%, D:1.9%] |
|  | F:1.7%, M:80.0%, n:1440 | F:3.0%, M:54.4%, n:1440 | F:0.7%, M:82.1%, n:1440 |
| **Transrate predicted ORFs** | 4,628 | 12,439 | 7,593 |
