## Supplementary figures and images for "Utilization of tissue ploidy level variation in *de novo* transcriptome assembly of *Pinus sylvestris*"

### Supplementary file 7

#### Slide 1
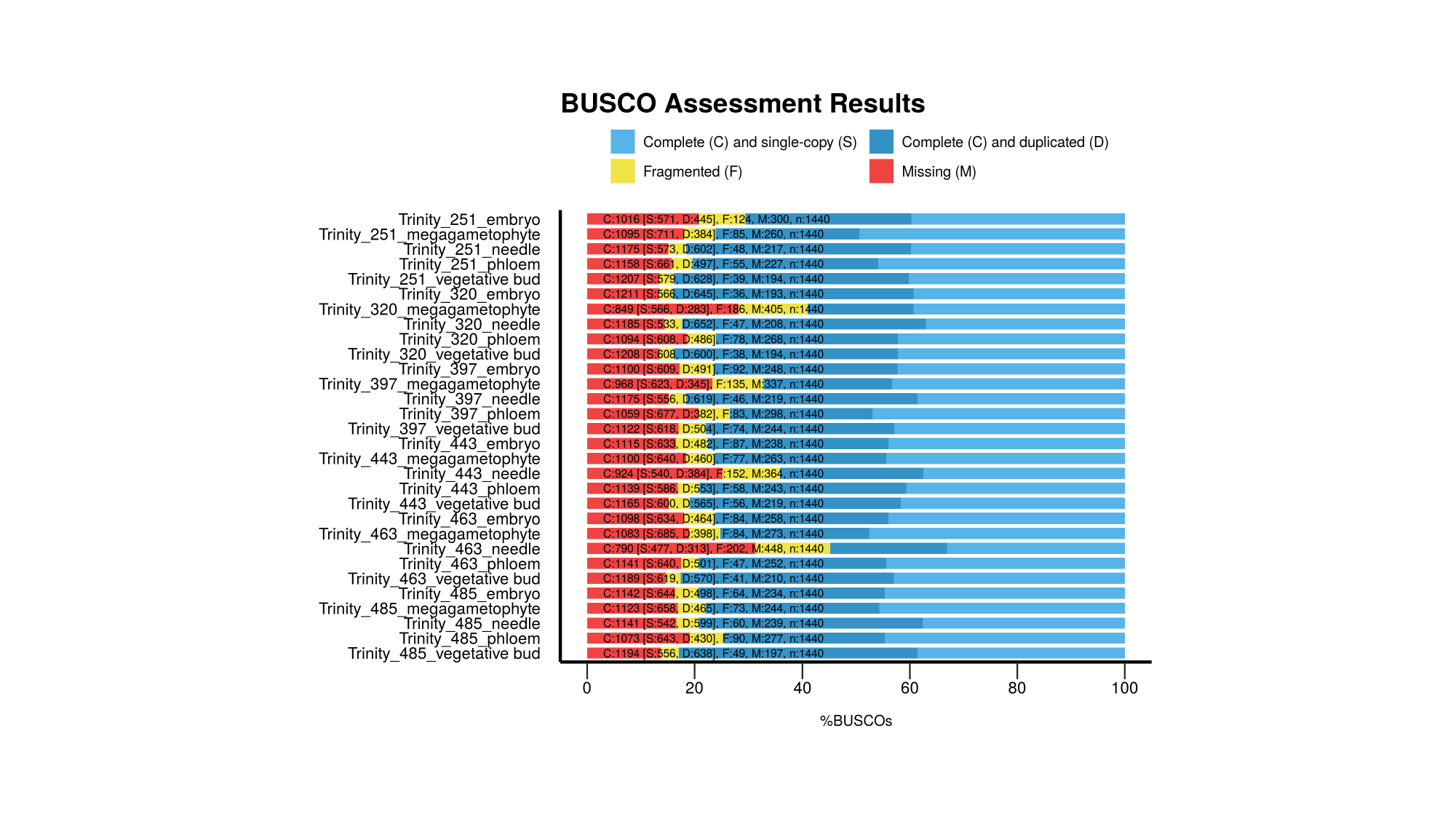

### Supplementary file 8

#### Slide 1
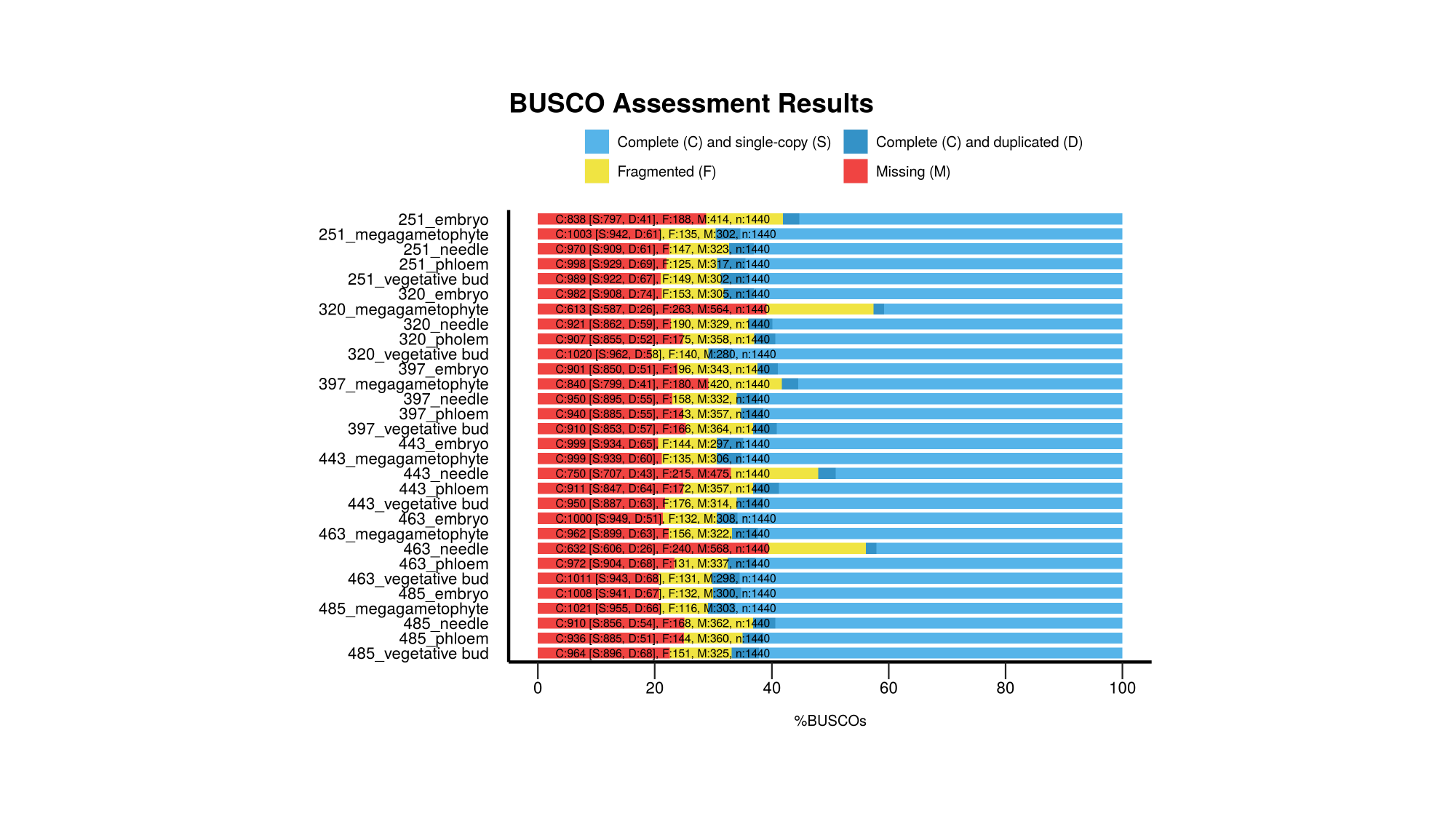

### Supplementary file 10

#### Slide 1
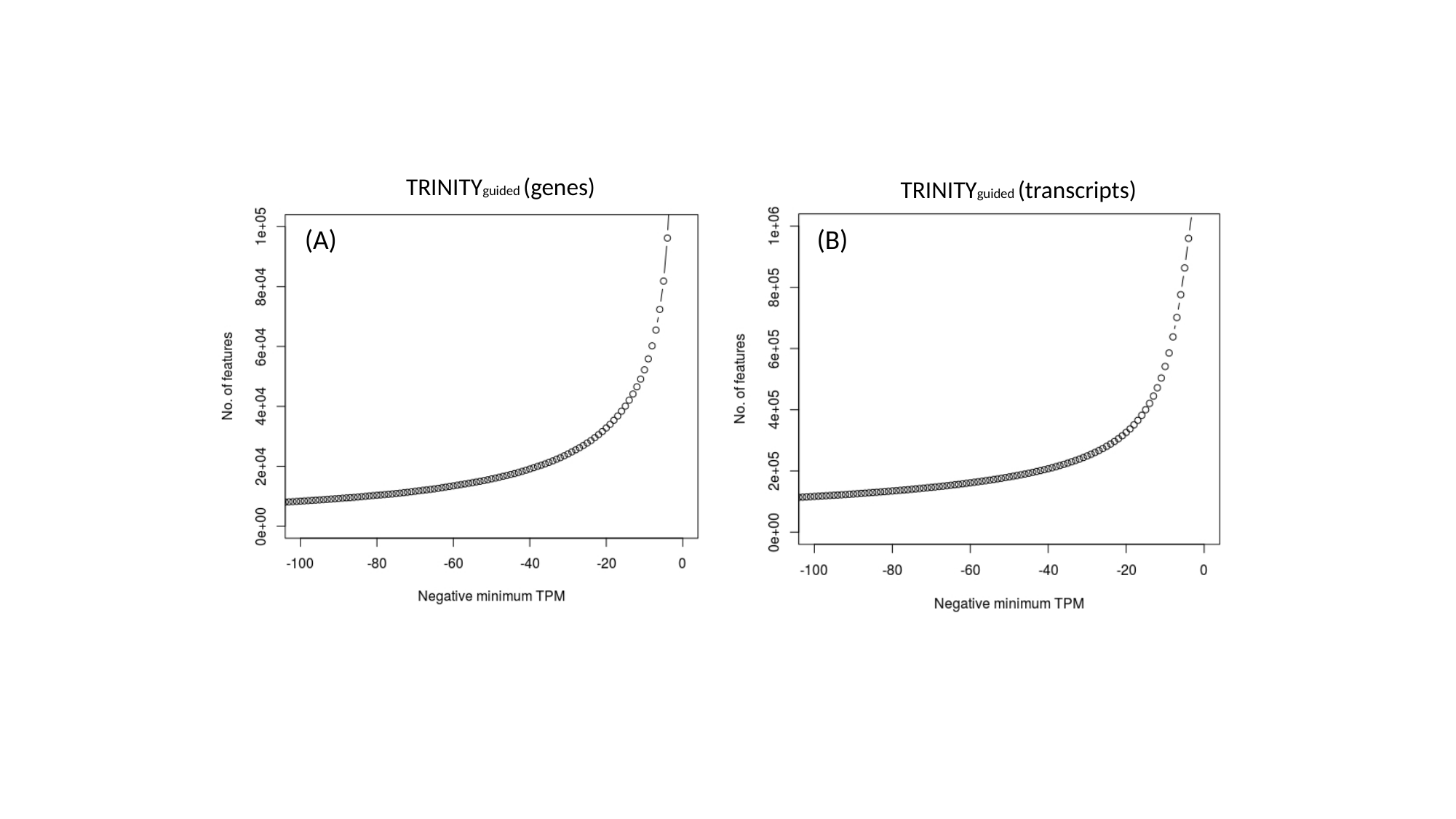

TRINITYguided (genes)
TRINITYguided (transcripts)
(A)
(B)

### Supplementary file 12

#### Slide 1
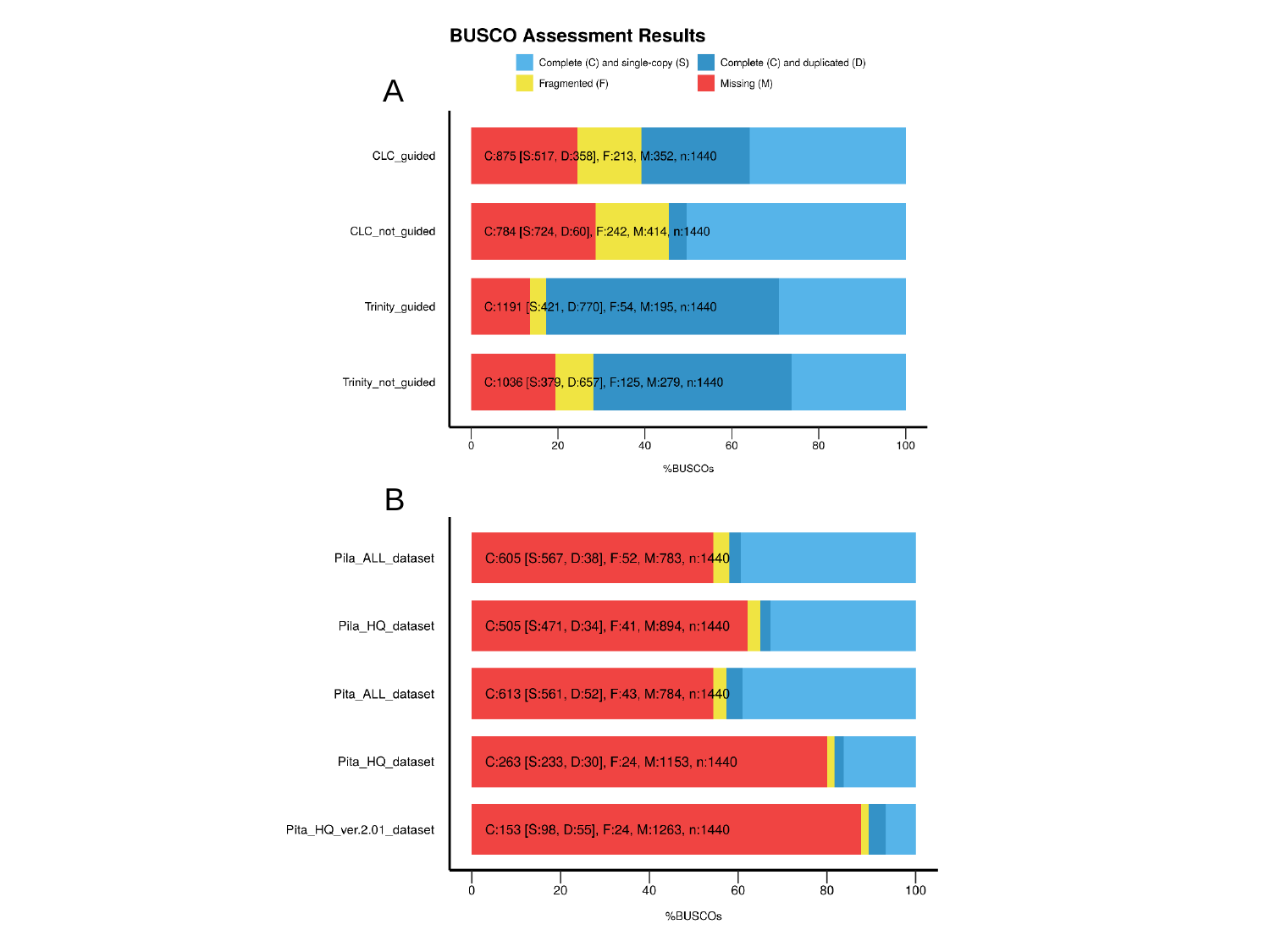

A
B
