## Supplementary material for "Utilization of tissue ploidy level variation in *de novo* transcriptome assembly of *Pinus sylvestris*"

### Slide 1
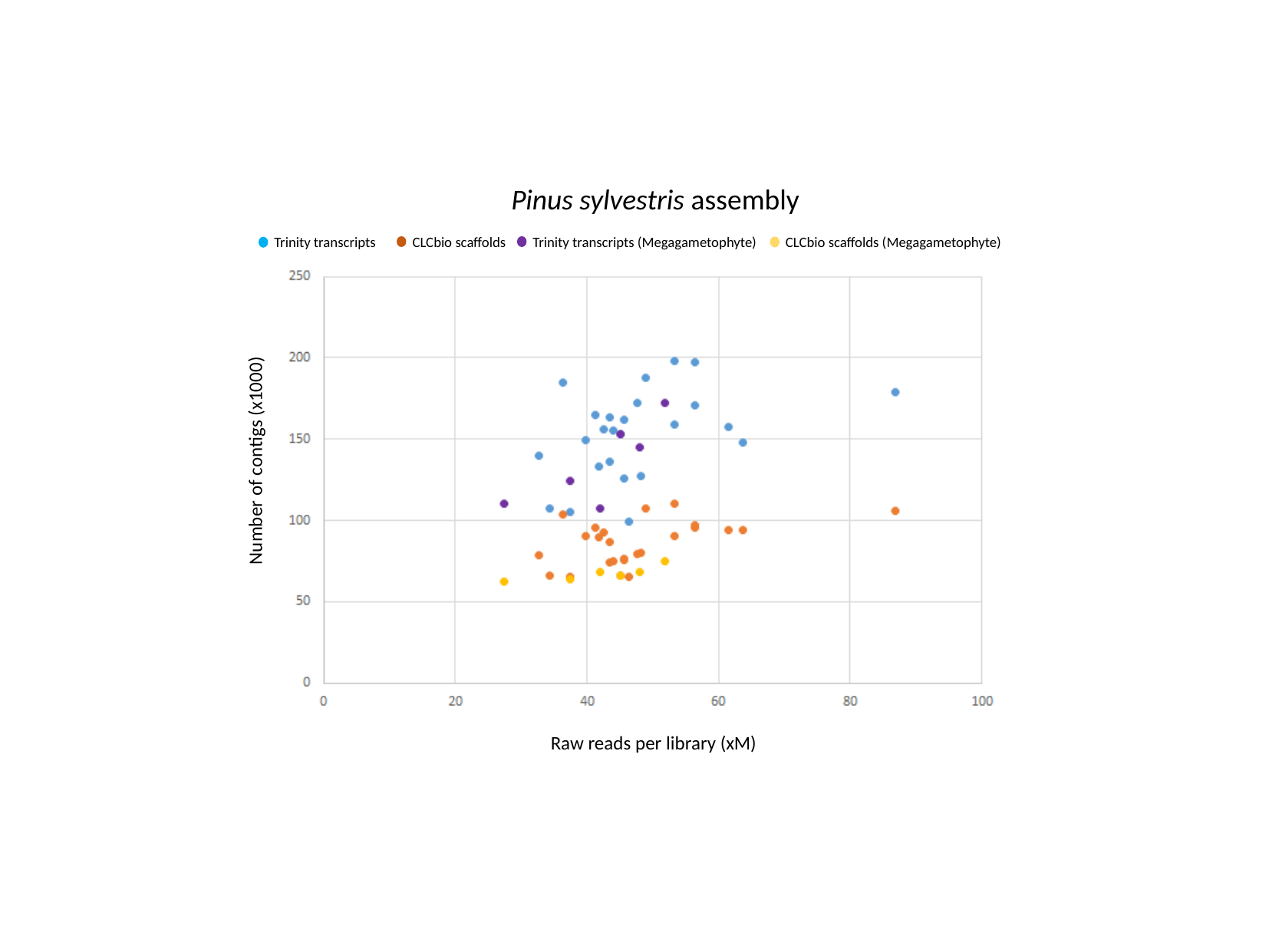

Pinus sylvestris assembly
CLCbio scaffolds
Trinity transcripts (Megagametophyte)
CLCbio scaffolds (Megagametophyte)
Trinity transcripts
Number of contigs (x1000)
Raw reads per library (xM)
