## Supplementary material for "Utilization of tissue ploidy level variation in *de novo* transcriptome assembly of *Pinus sylvestris*"

### Slide 1
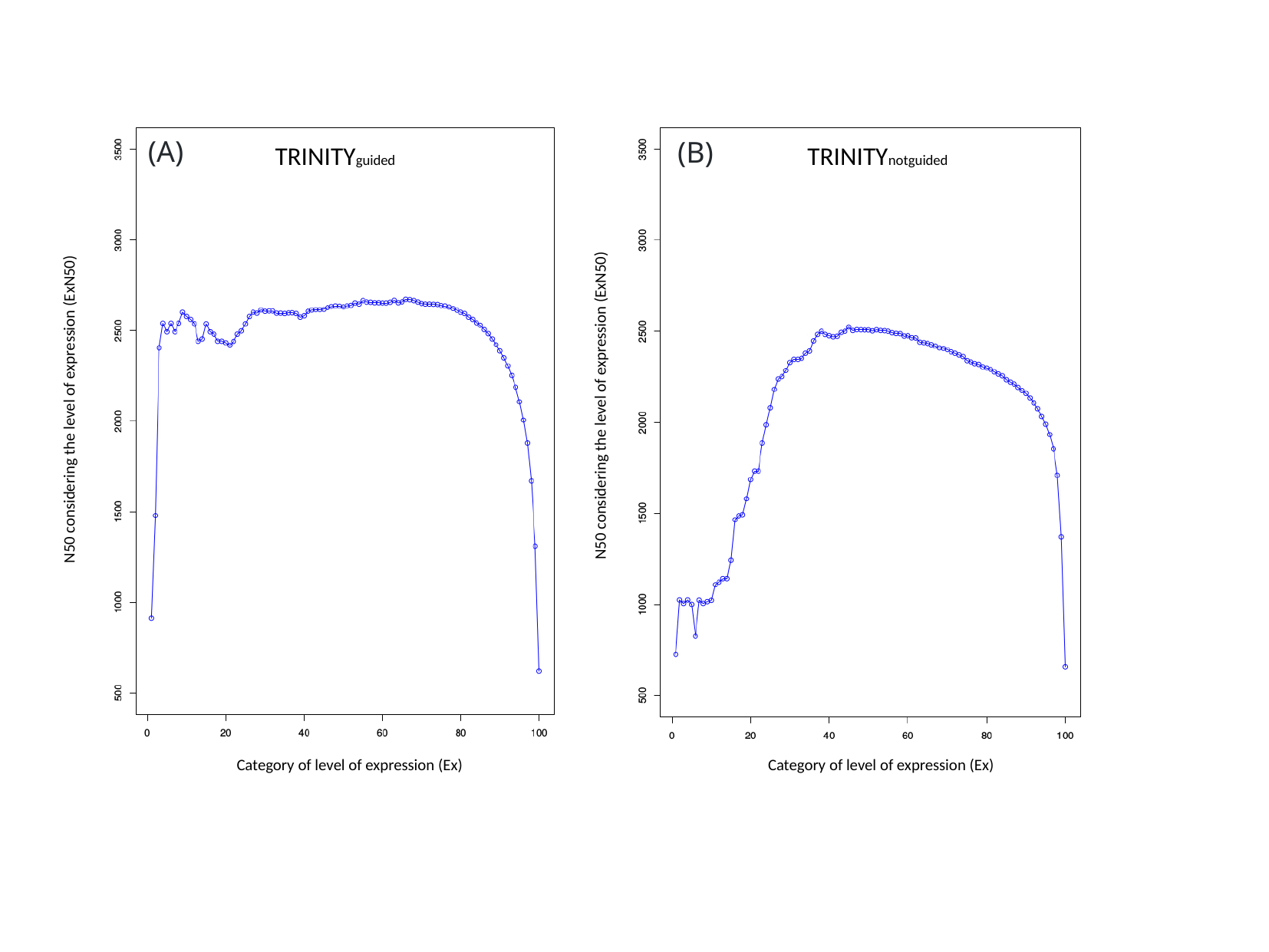

(A)
(B)
TRINITYnotguided
TRINITYguided
N50 considering the level of expression (ExN50)
N50 considering the level of expression (ExN50)
Category of level of expression (Ex)
Category of level of expression (Ex)
