## Supplementary material for "Utilization of tissue ploidy level variation in *de novo* transcriptome assembly of *Pinus sylvestris*"

### Slide 1
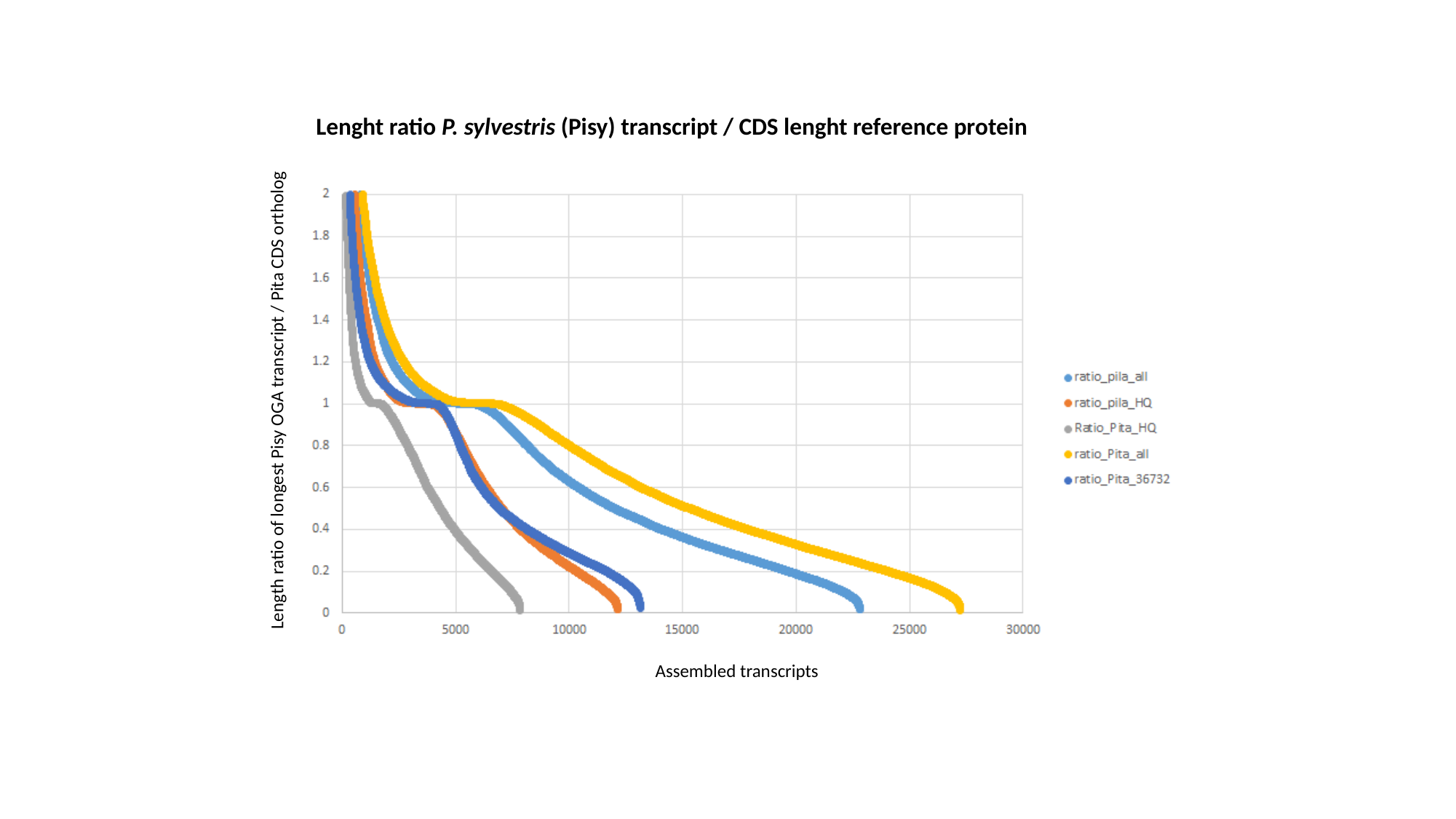

Lenght ratio P. sylvestris (Pisy) transcript / CDS lenght reference protein
Length ratio of longest Pisy OGA transcript / Pita CDS ortholog
Assembled transcripts
